## Supplemental Figures and Table Descriptions for "Transcription factor regulation of eQTL activity across individuals and tissues"

### Supplementary Information for *Transcription factor regulation of eQTL activity across individuals and tissues*

- Supplementary Figures 1-17
- Supplementary Table legends
- Supplementary Tables 1-5 available in Excel file

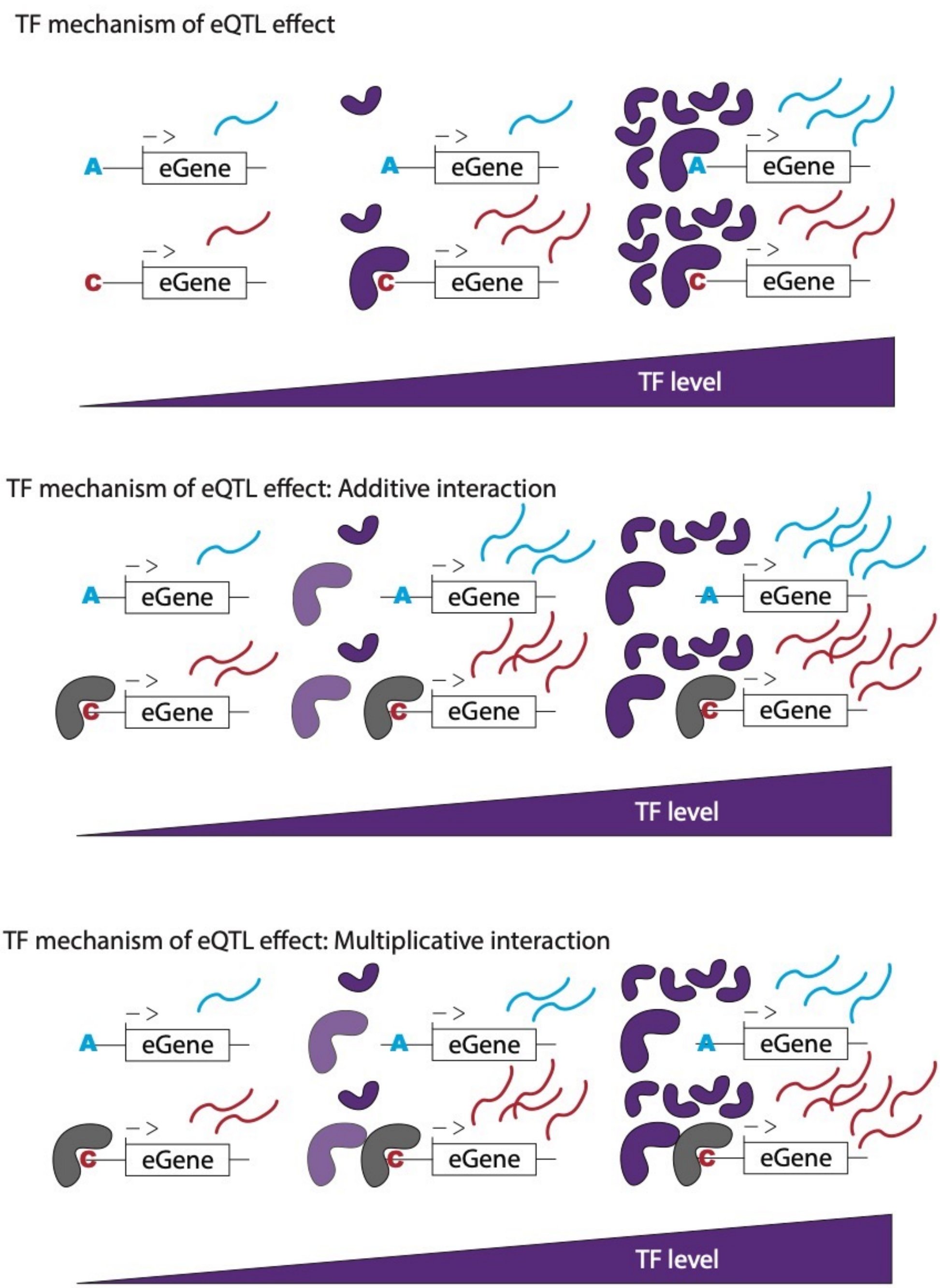


**SFigure 1. Possible models of TF mechanisms of eQTL effect variability.** (Top) An eQTL variant increases the affinity of the purple TF, resulting in observable eQTL effects at mid-range levels of purple TF. (Middle) An eQTL variant disrupts the affinity of the gray TF, while the purple TF binds to a different region of the *cis*-regulatory region which interacts additively with the region where the gray TF binds. The eQTL effect will be observable at low purple TF levels, but may become overpowered and unobservable if purple TF levels increase and expression generated by the purple TF locus exceeds that generated by the gray TF locus. (Bottom) An eQTL variant disrupts the affinity of the gray TF, while the purple TF binds binds to the gene's *cis*-regulatory region and interacts multiplicatively with the region where the gray TF binds. The eQTL effect should remain constant at all purple TF levels.

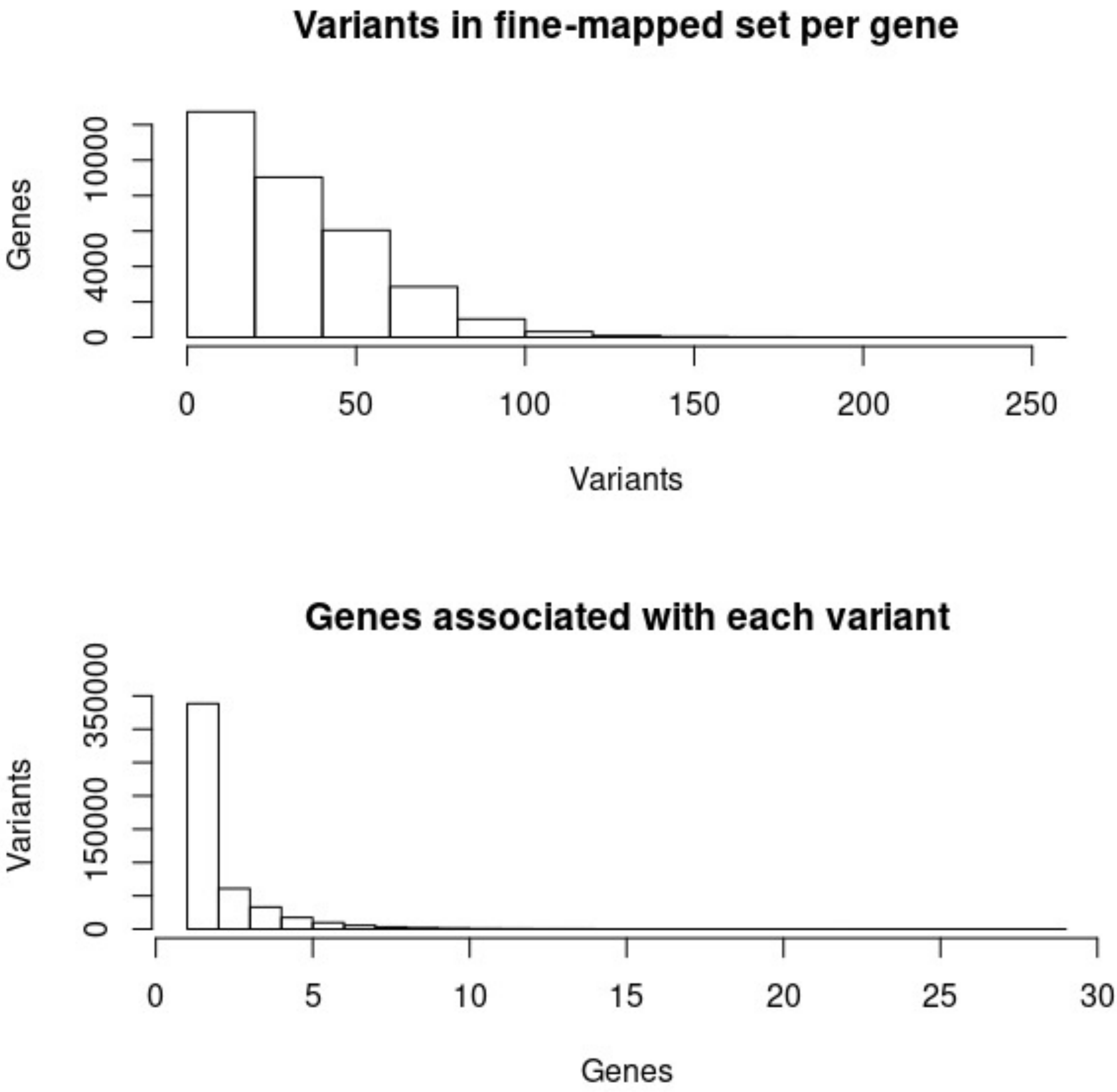


**SFigure 2. Variant-gene associations of potentially causal eQTL variants.** Caviar fine-mapped variants were overlapped with TF ChIPseq and motif variants. The number of resulting fine-mapped variants per gene (top) and genes per variant (bottom) are displayed.

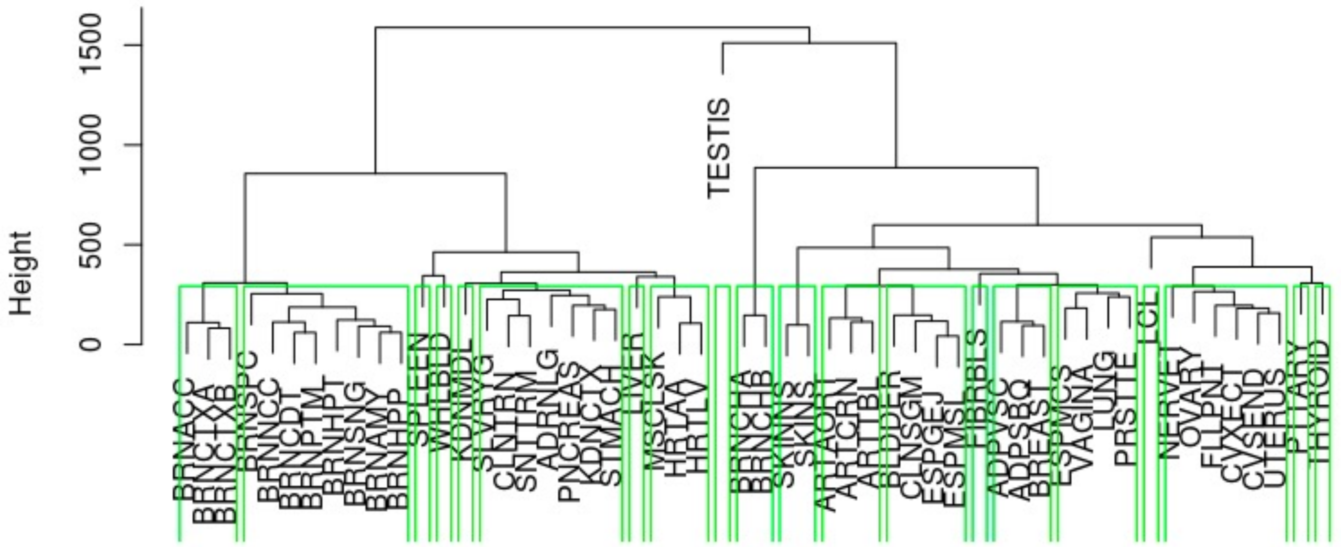


**SFigure 3. Clustering of GTEx tissues based on gene expression.** We clustered tissues based on median TPM across all genes using Euclidean distances and Ward.D clustering, cut the resulting tree to generate twenty clusters (green boxes). The tissue with the largest sample size was selected from each cluster.

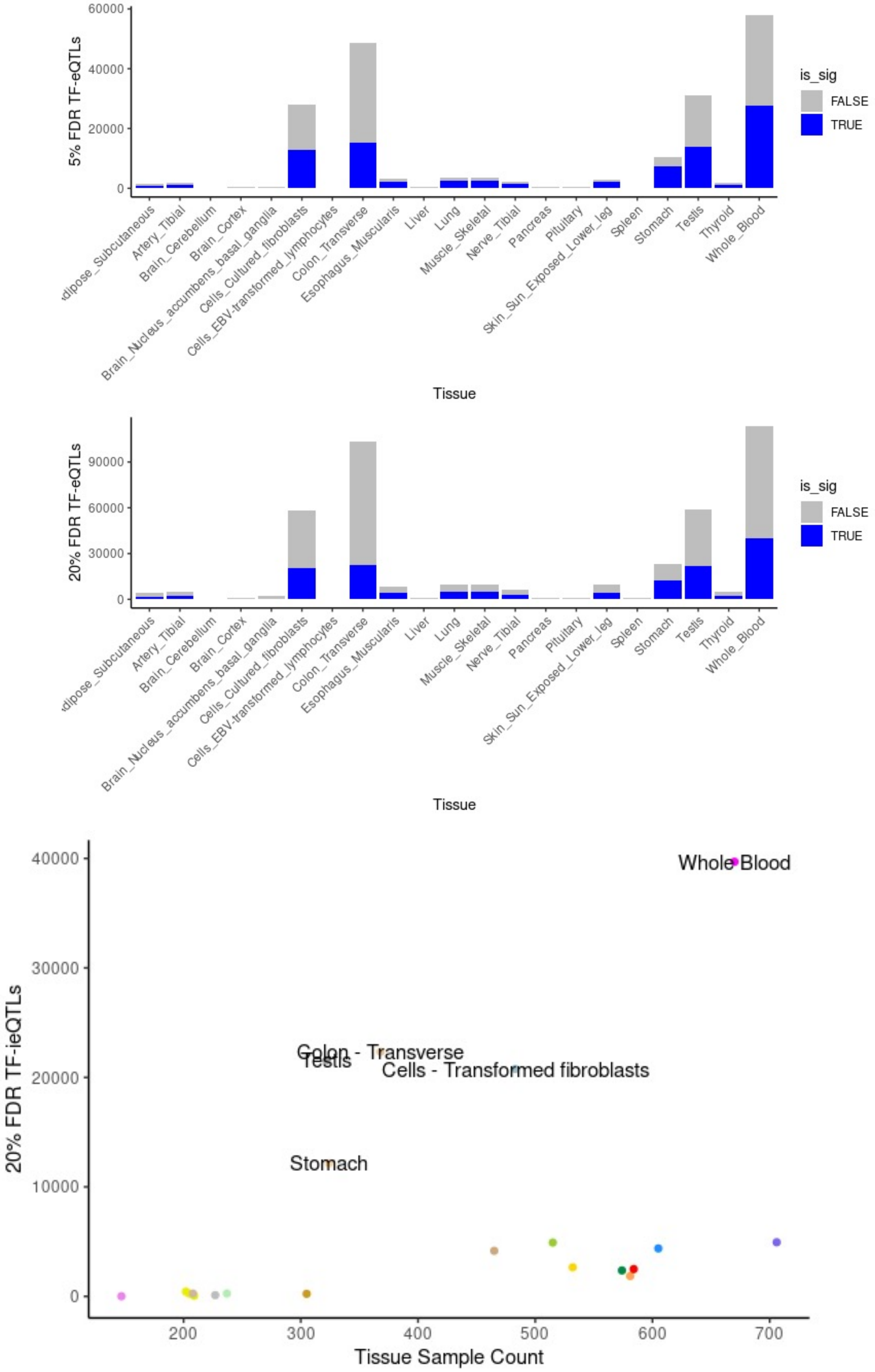


**SFigure 4. Within-tissue TF-eQTLs.** (Top) 5% and 20% FDR TF-eQTLs per tissue, colored by whether or not top TF-eQTL variant was significantly associated with gene expression in that tissue. Only TF-eQTL variants with a significant eQTL were retained for further analysis. Since we performed additional filtering of these hits, we chose a less stringent FDR (20%) for further analysis. (Bottom) Tissues plotted by number of TF-eQTLs vs. tissue sample size. Outlier tissues are labeled.

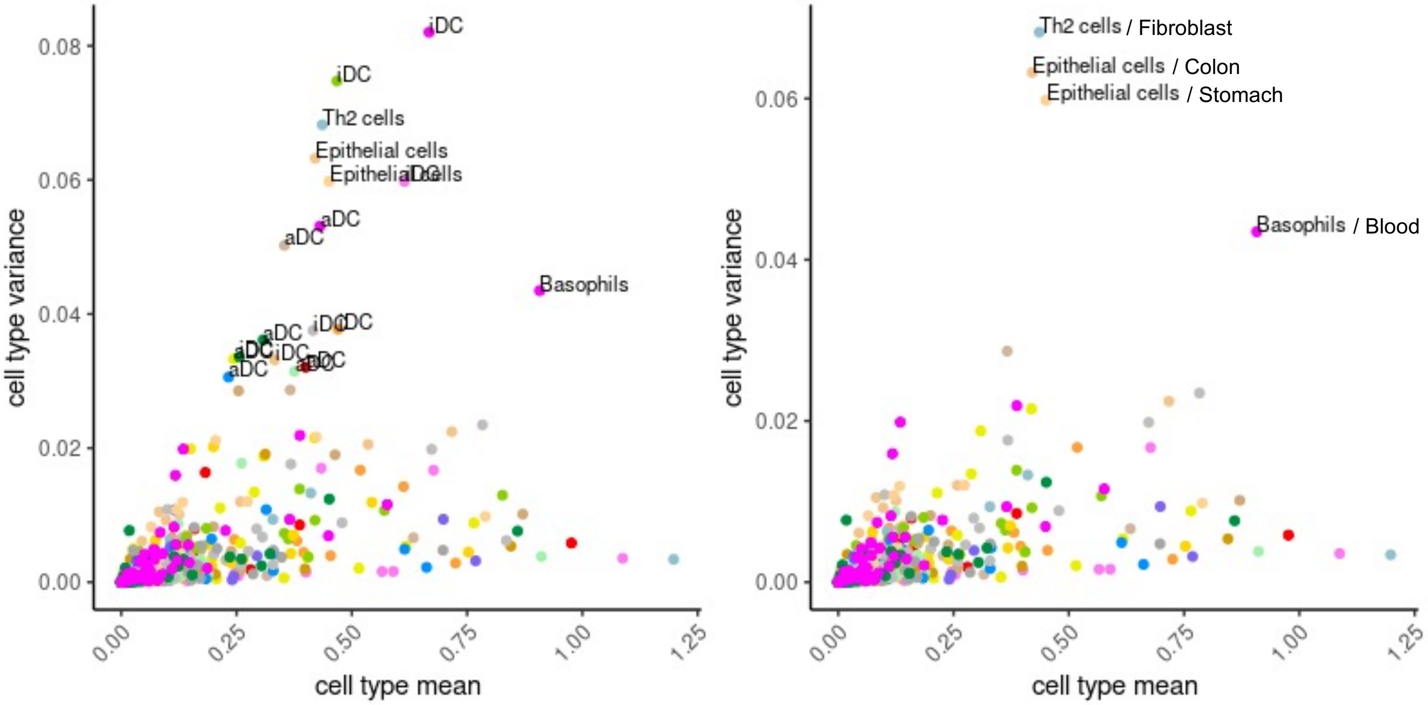


**SFigure 5. Cell type composition variability of GTEx tissues.** Cell type enrichments were calculated *in silico* using XCell, and the mean/variance of each cell type in each tissue was calculated. Both plots show cell type variance vs. mean per tissue (dot color). aDC and iDC estimates frequently had large variance (left), thus they were removed (right). Four tissues remained with large cell type variance: fibroblasts, colon, stomach, and blood.

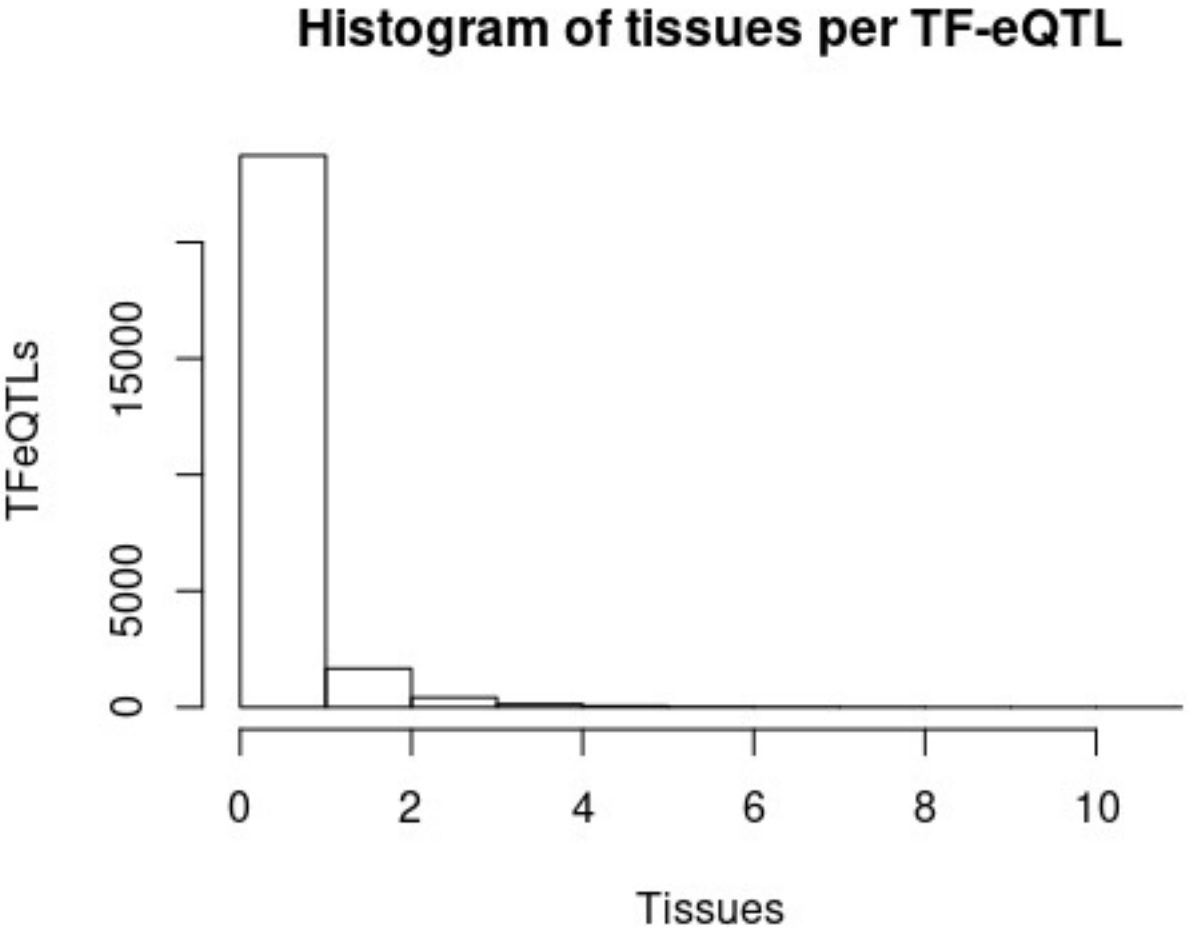


**SFigure 6. Histogram of 20% FDR TF-eQTLs across tissues.** The majority of TF-eQTLs were seen in one tissue only, though 2315/26044 (8.9%) of TF-eQTLs were observed in more than one tissue.

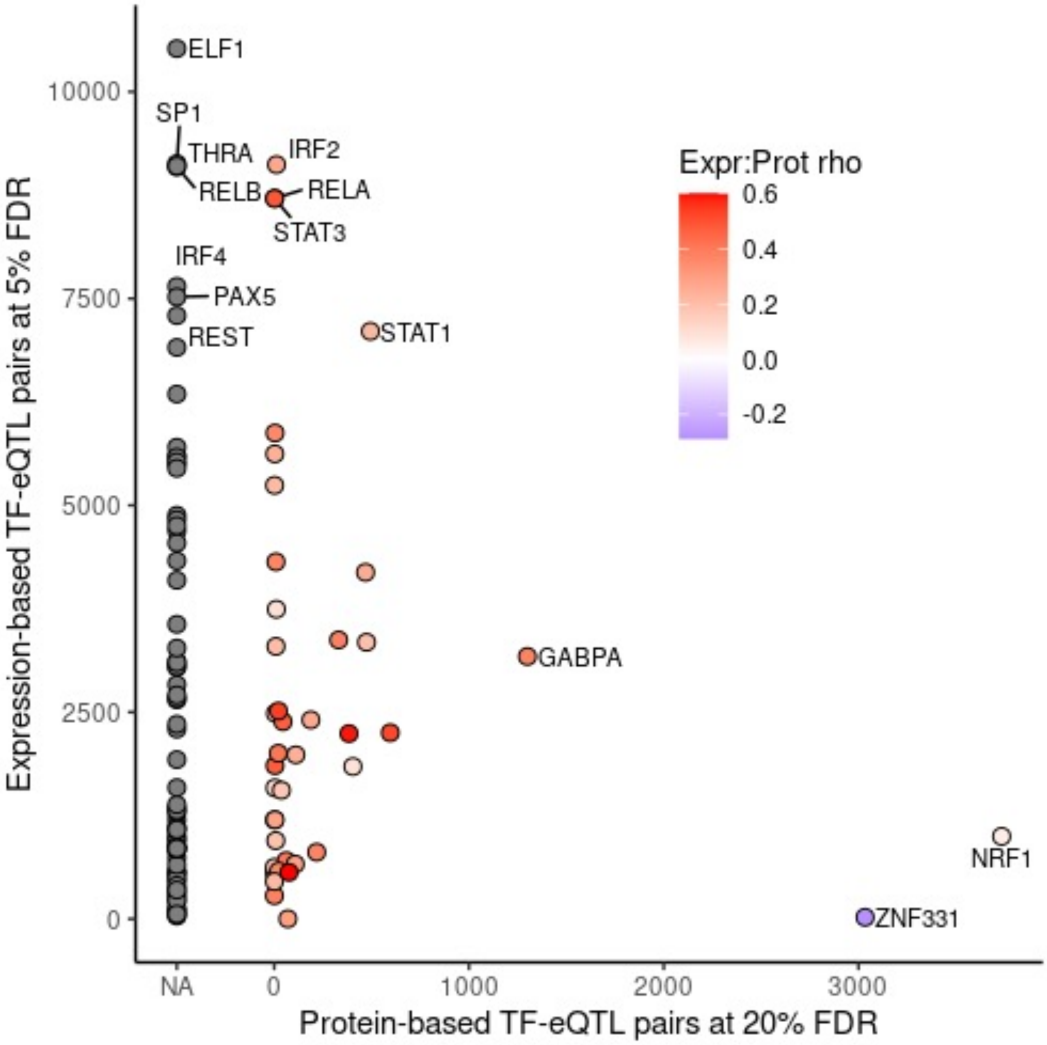


**SFigure 7. Cross-tissue TF-eQTLs per TF.** Number of cross-tissue expression-based TF-eQTLs are plotted versus number of cross-tissue protein-based TF-eQTLs. Dots are colored by Spearman correlation of cross-tissue median TF protein and expression levels.

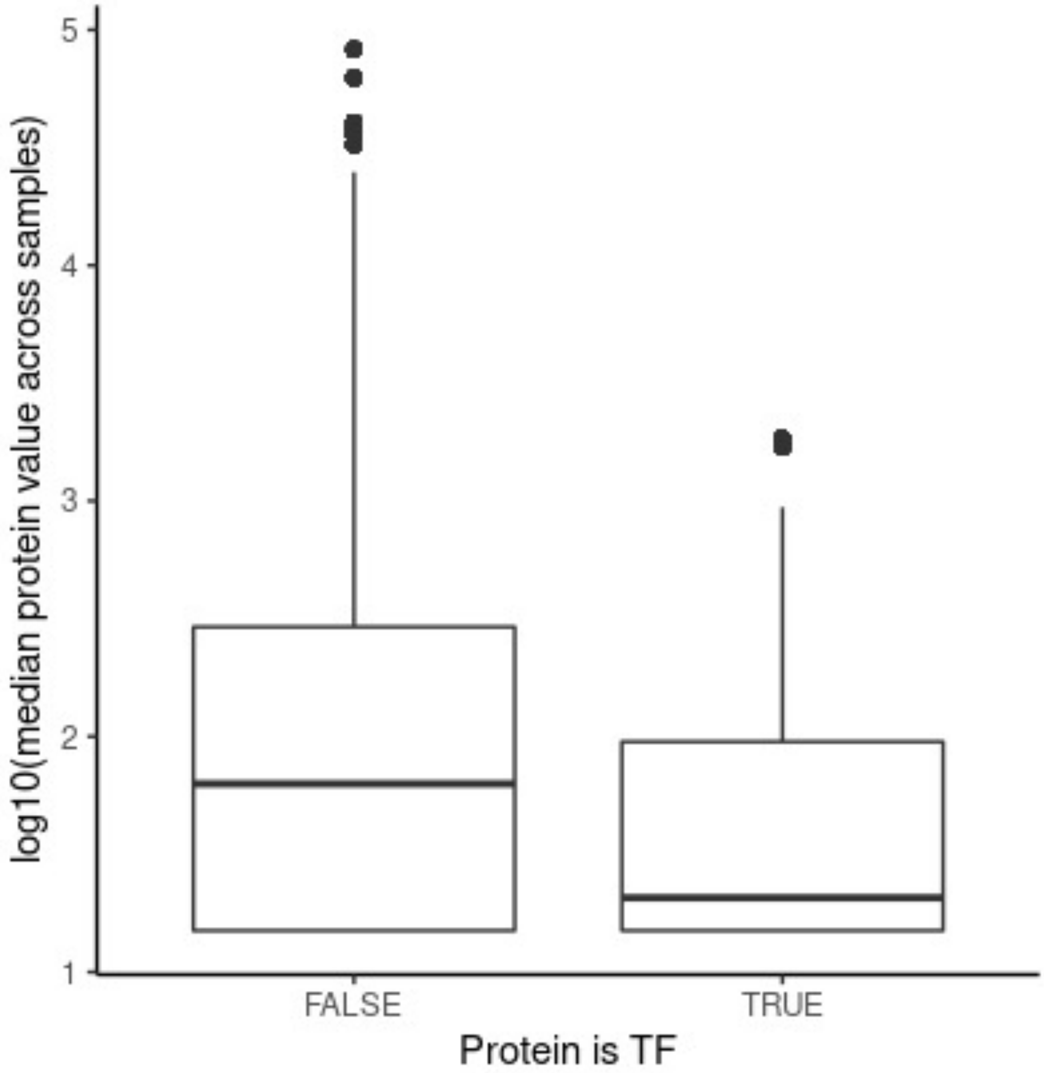


**SFigure 8. TF protein levels.** Log10 of median relative protein abundance for all measured proteins, subset by whether the protein is a tested TF or not.

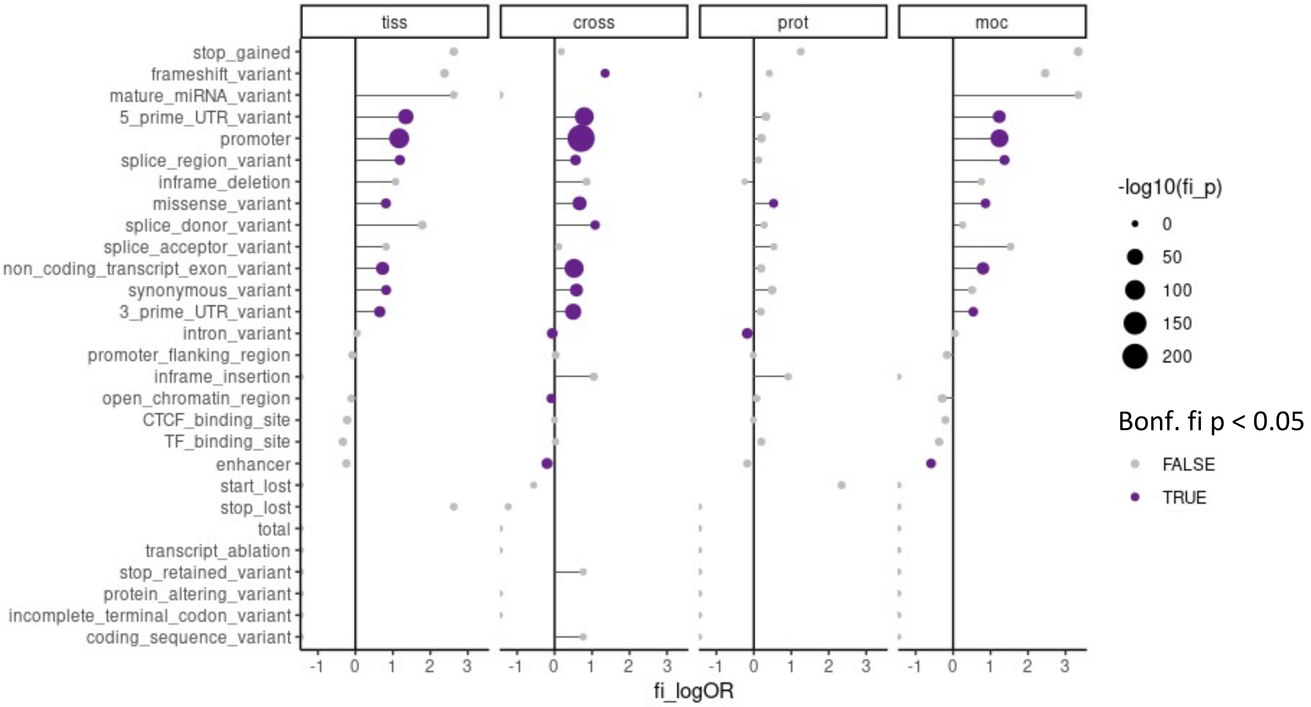


**SFigure 9. Annotation overlap of top TF-eQTL variants.** Enrichment of top TF-eQTL variant in each dataset for various genomic annotations from Ensembl Variant Effect Predictor.

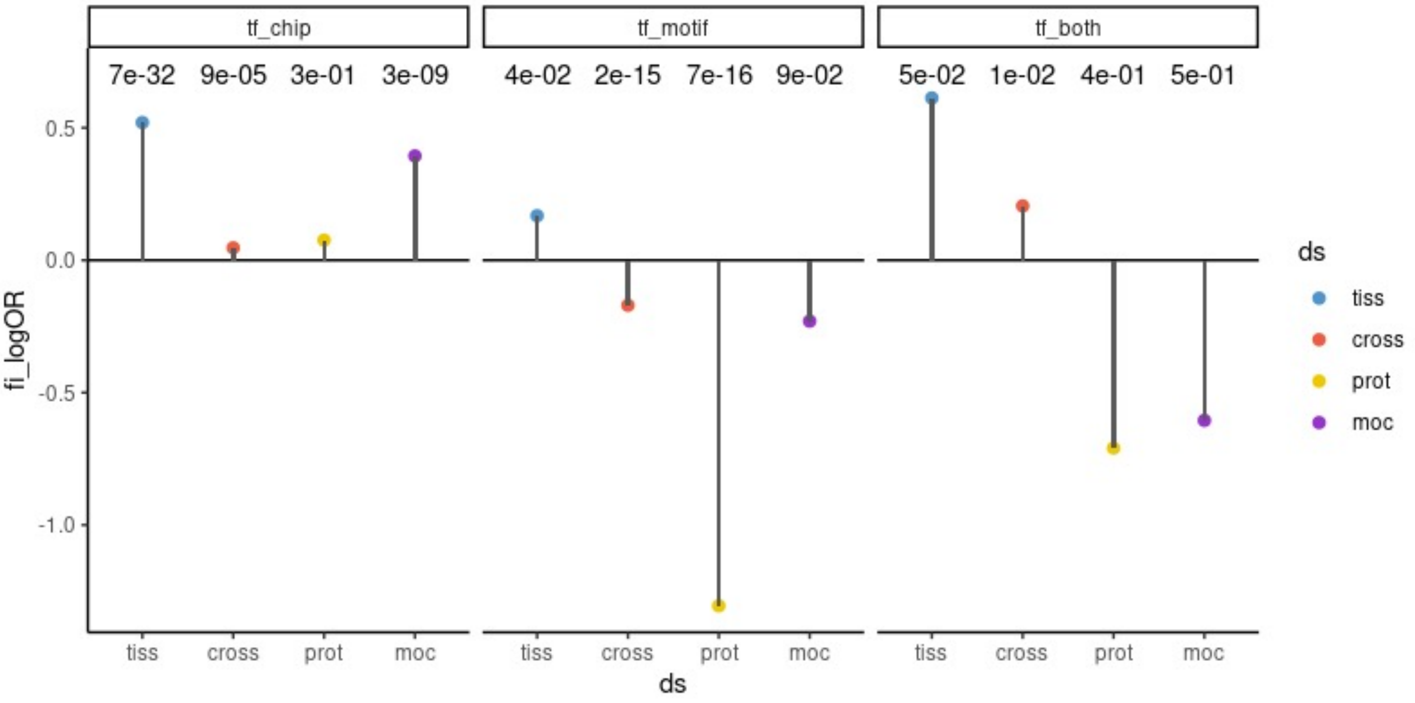


**SFigure 10. TFBS overlap of top TF-eQTL variants.** Enrichment of top TF-eQTL variant in each dataset for TF overlap, as defined by ENCODE TF ChIPseq peaks, HOCOMOCO predicted motifs, or both annotations together. Fisher's exact test p-values are plotted.

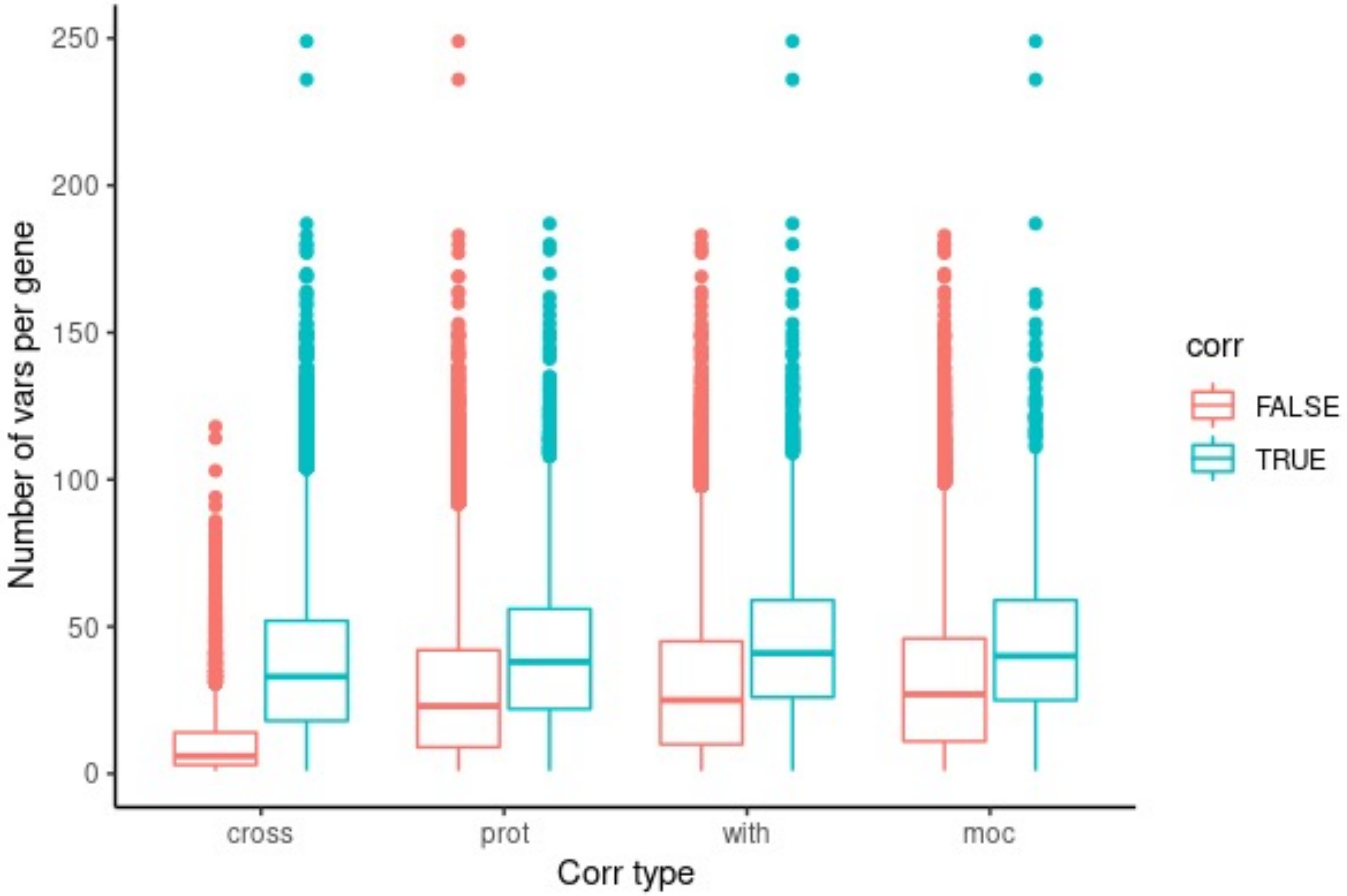


**SFigure 10. Tested variants per gene.** Number of tested variants per gene, separated by whether the gene had a TF-eQTL (color) in the given dataset (x-axis). Significant TF-eQTL genes tended to have more tested variants per gene than unsignificant genes.

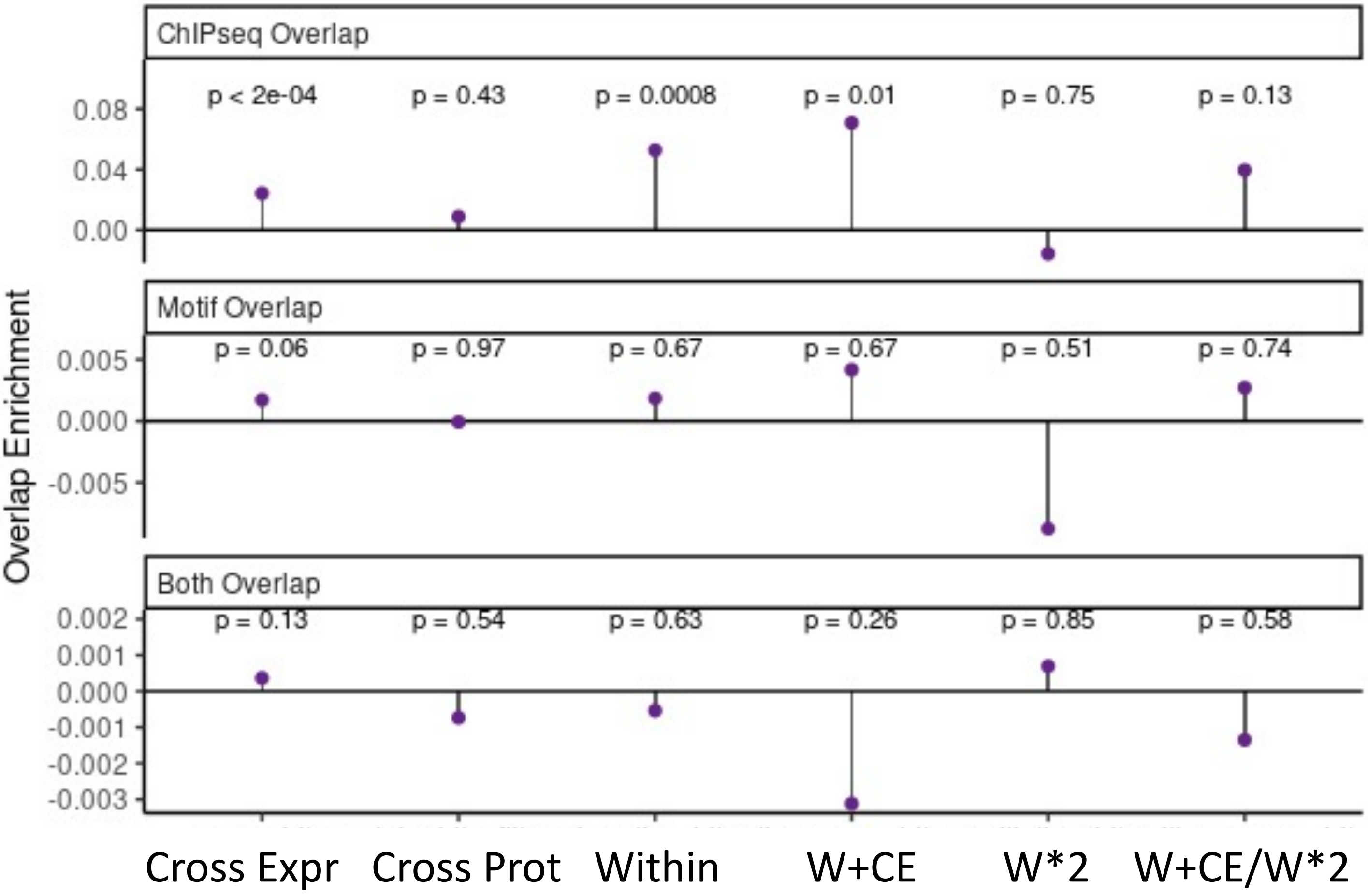


**SFigure 11. TF binding overlap enrichment.** Overlap enrichment for each TF-eQTL dataset for ChIPseq overlap, motif overlap, and both overlap.

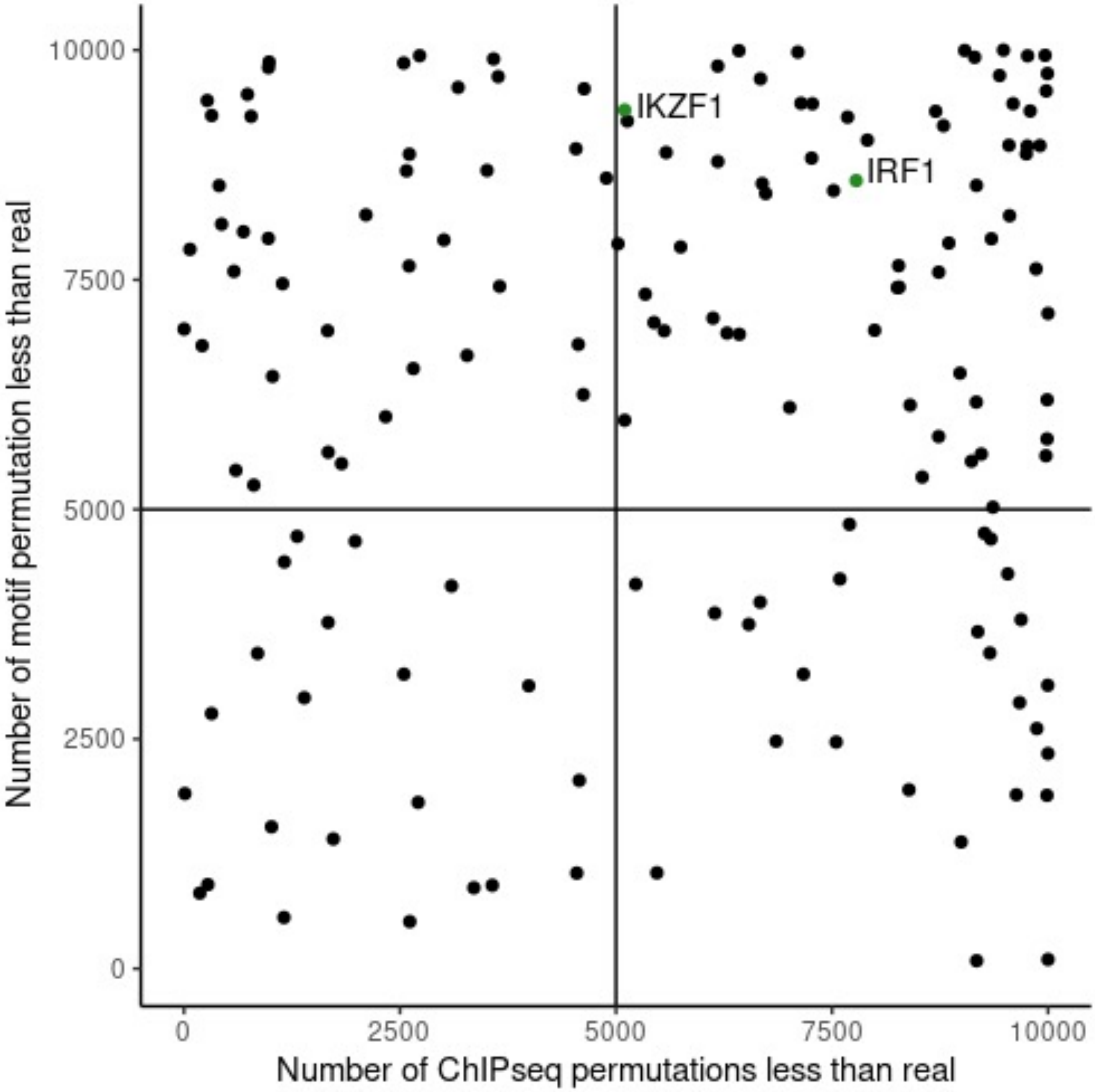


**SFigure 12. TF binding overlap enrichment permutations.** Number of permuted datasets with an enrichment statistic lower than the real dataset, plotted per TF.

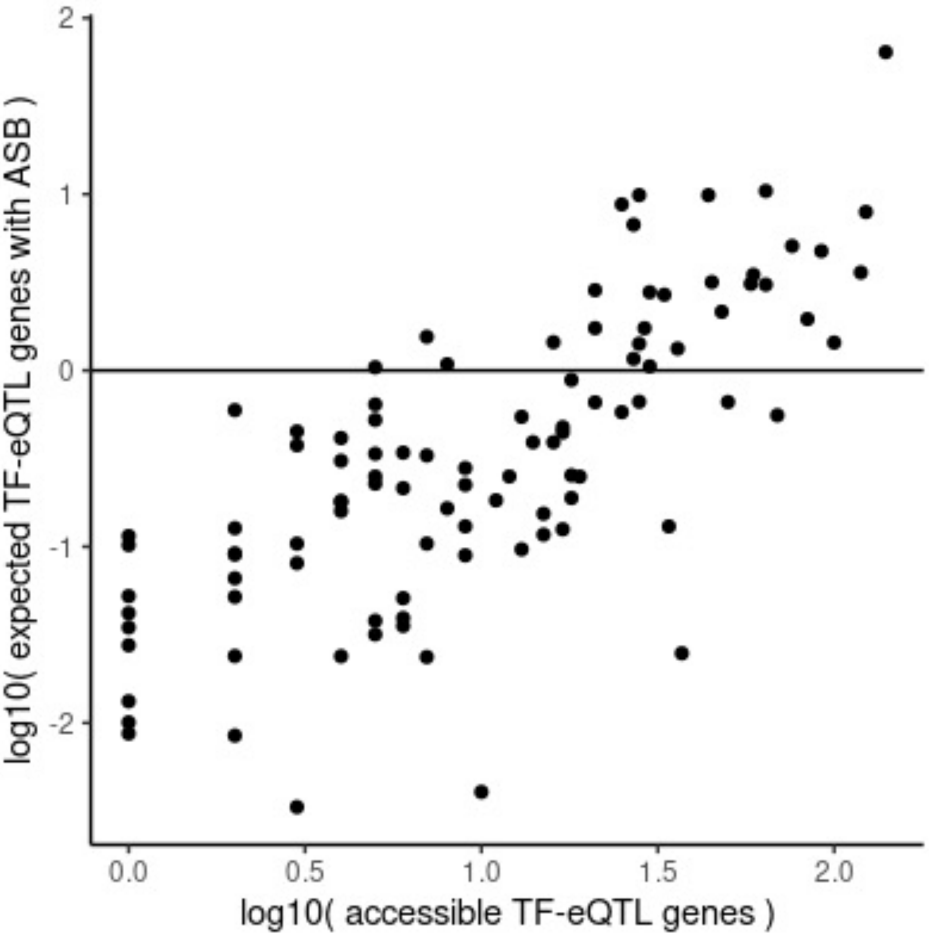


**SFigure 13. Expected ASB overlap.** Number of expected TF-eQTL genes with ASB calculated by number of TF-eQTL gene variants accessible for a TF times percent of all accessible variants with ASB for that TF.

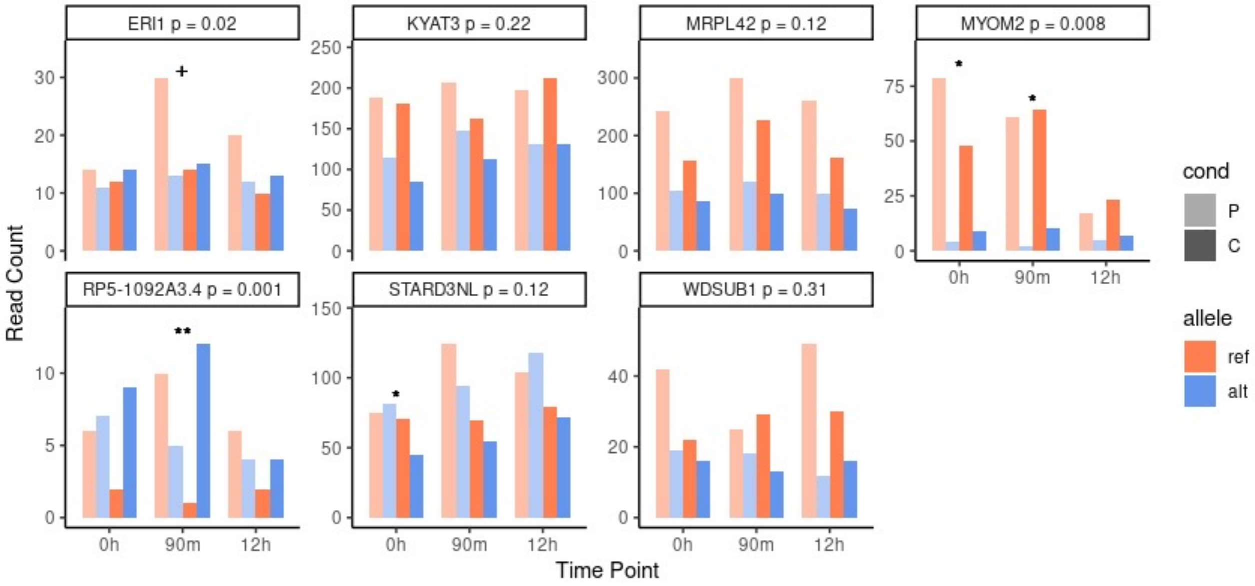


**SFigure 14. ASE in HEK293T cells for dual-evidence TF-eQTL genes.** Read counts are plotted for three timepoints relative to LPS immune stimulation. Fisher's exact test p value for data combined across timepoints is displayed at the top of the chart, while significance for individual timepoints is denoted by symbols: + p<0.10, * p<0.05, ** p<0.01.

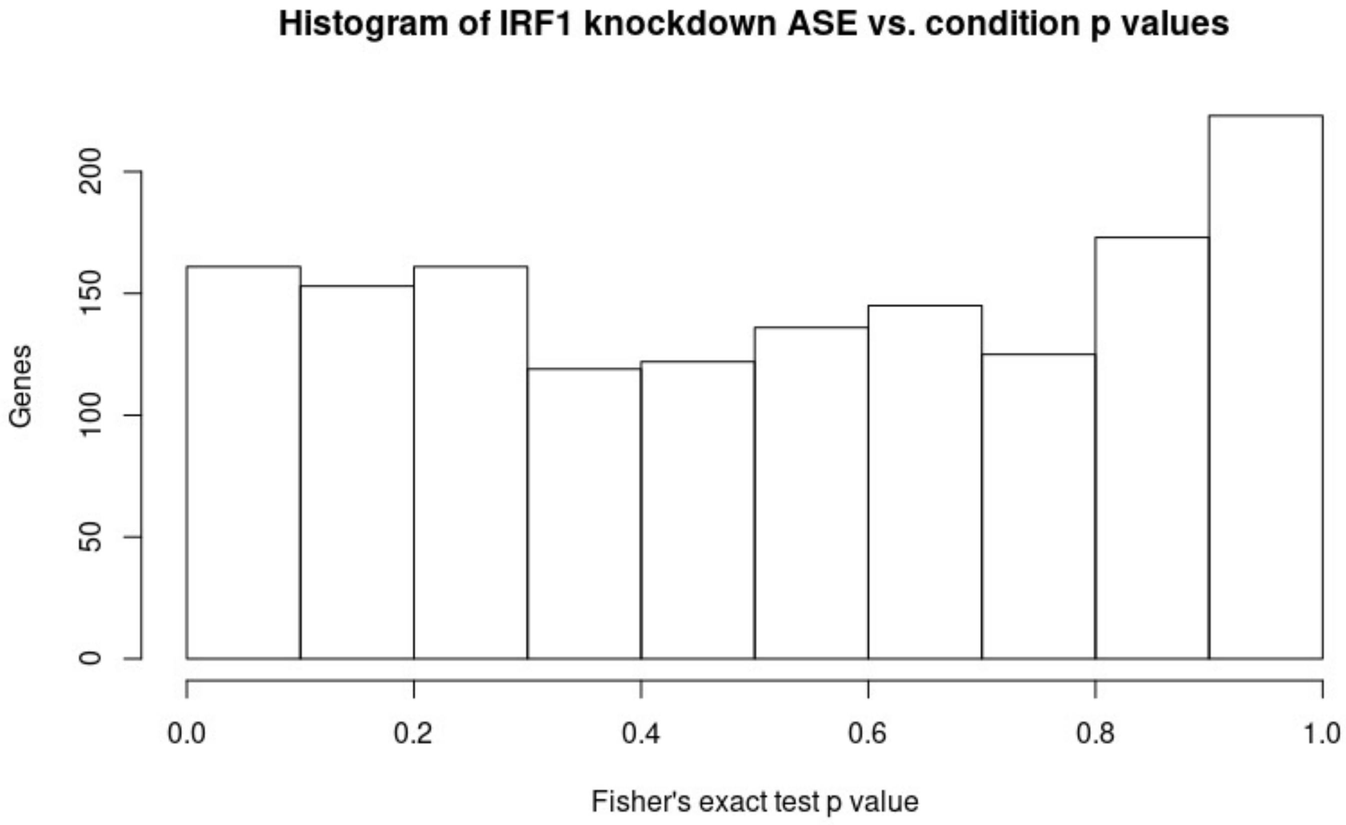


**SFigure 15. P values of IRF1 knockdown effects on ASE.** Fisher's exact tests were run on 2x2 tables of allelic read counts in IRF1 knockdown and control experiments for all adequately covered genes.

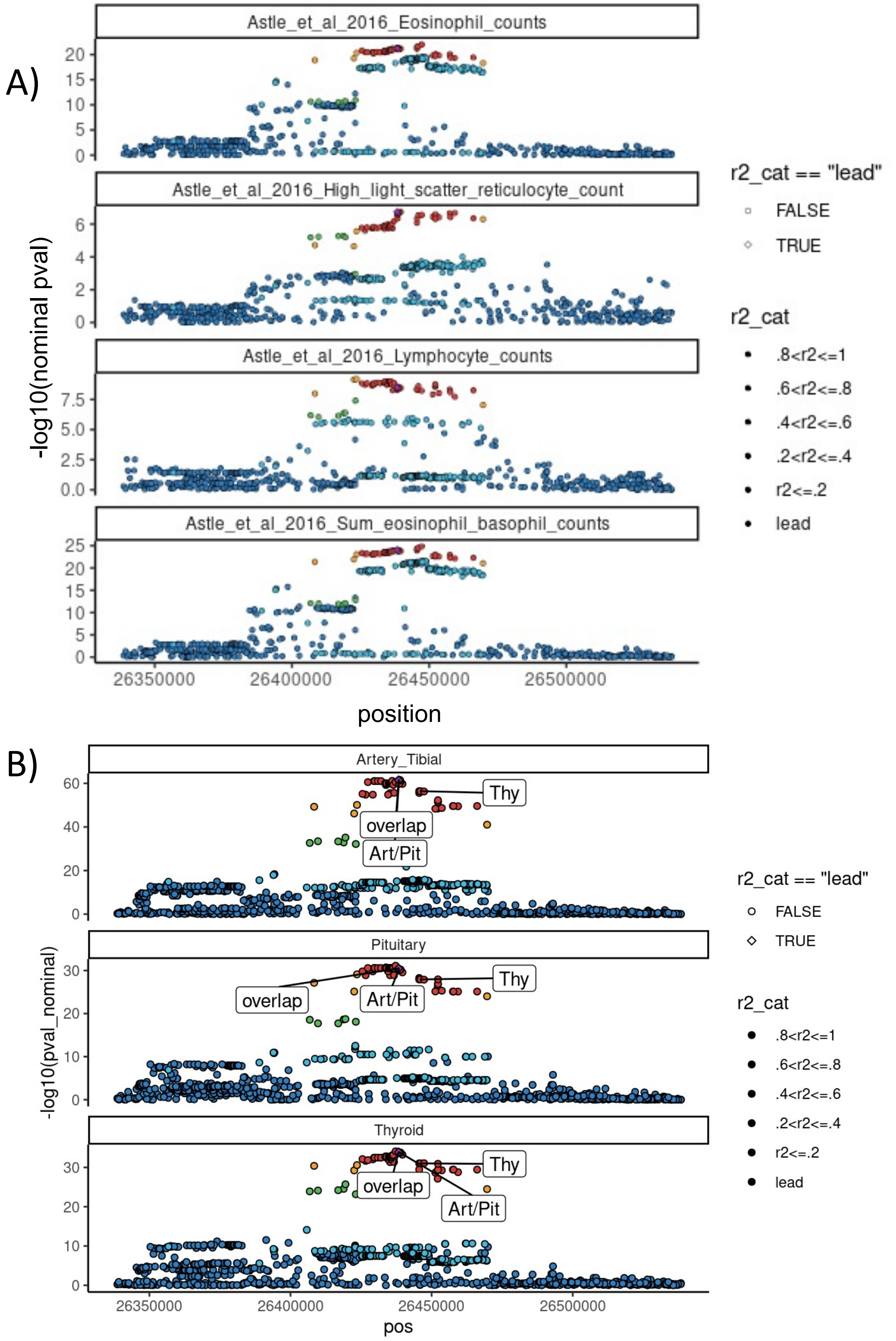


**SFigure 16. LocusZoom plots of GWAS and eQTL.** A) P values are plotted for four blood cell GWAS traits that colocalize with an *APBB1IP* eQTL in any tissue. B) P values are plotted for *APBB1IP* eQTL in three tissues with an IKZF1-eQTL signal. Top IKZF1-eQTL variants are labeled, as well as a variant that overlaps an IKZF1 motif and ChIP-seq peak (overlap).

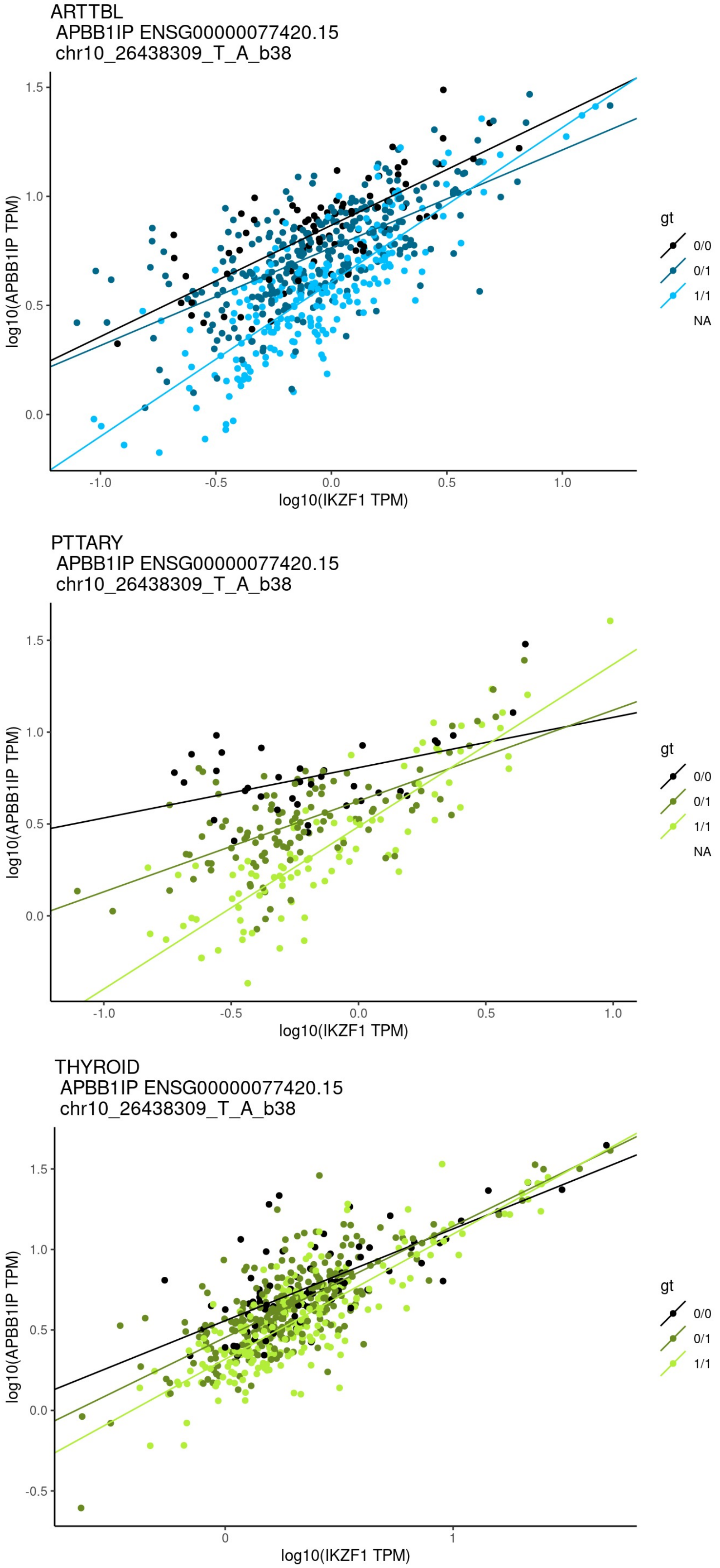


**SFigure 17. IKZF1-eQTL interaction signals.** Individuals are plotted by *APBB1IP* expression vs. IKZF1 TF expression in three tissues. Trend lines show linear regression lines per genotype. All three tissues had a significant IKZF1-eQTL.

IKZF1 motif logo

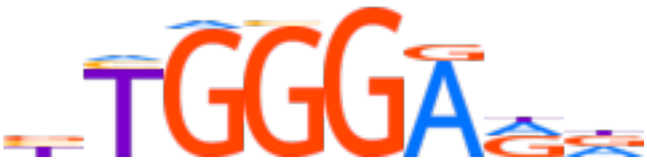

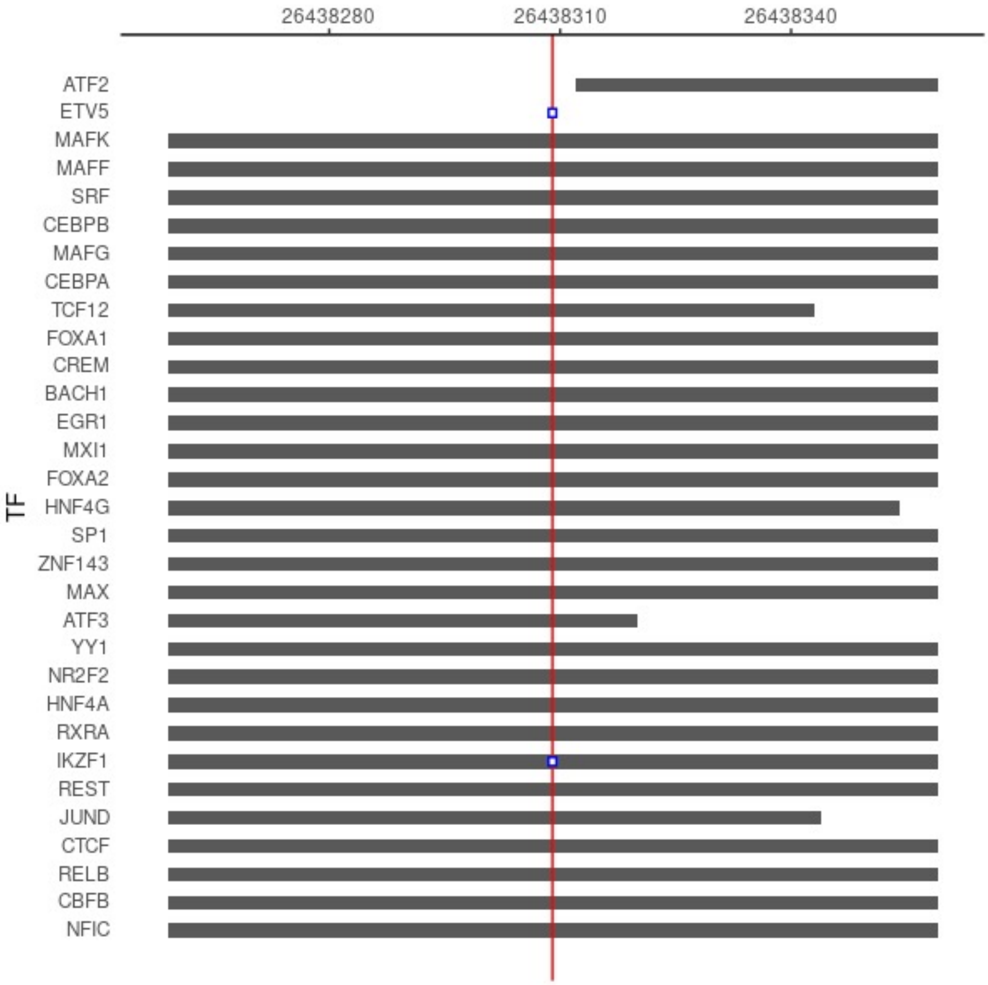


**SFigure 18. TF binding information of *rs1335540*.** rs1335540 (red line) overlaps multiple TF ChIPseq peaks and two TF motifs. IKZF1 is the only TF which the SNP overlaps both of. The IKZF1 motif logo is displayed at the top of the figure.

### Supplementary Table legends

STable 1. GTEx Tissues. Information on available expression and protein samples for GTEx tissues.

STable 2. Dual Evidence TF-eQTLs. TF-eQTL genes with two lines of supporting evidence (2+ tissues or cross + within tissue). Each row represents a TF-eQTL in a single dataset.

STable 3. GxE Overlap. Overlap of dual evidence TF-eQTL genes with gene-by-environment interacting genes from Findley et al., 2021.

STable 4. GWAS Coloc TF-eQTLs. eQTL genes in tissues that both colocalize with a GWAS trait and have a dual evidence TF-eQTL in that tissue, filtered for $r^2 > 0.4$ between the lead TF-eQTL variant and lead colocalizing variant.

STable 5. Colocalizations between *APBB1IP* eQTLs and GWAS traits.
